# Vision is not olfaction: impact on the insect Mushroom Bodies connectivity

**DOI:** 10.1101/2024.08.31.610627

**Authors:** Florent Le Moël, Antoine Wystrach

**Affiliations:** School of Informatics, University of Edinburgh, Edinburgh EH8 9AB, UK; Centre de Recherches sur la Cognition Animale, CNRS, Université Paul Sabatier, Toulouse 31062 Cedex 09, France

**Keywords:** desert ants, mushroom bodies, visual-encoding, navigation, exploration, modelling

## Abstract

The Mushroom Bodies, a prominent and evolutionary conserved structure of the insect brain, are known to be the support of olfactory memory. There is now evidence that this structure is also required for visual learning, but the hypotheses about how the view memories are encoded are still largely based on what is known of the encoding of olfactory information. The different processing steps happening along the several relays before the Mushroom Bodies is still unclear, and how the visual memories actually may allow navigation is entirely unknown. Existing models of visual learning in the Mushroom Bodies quickly fall short when used in a navigational context. We discuss how the visual world differs from the olfactory world and what processing steps are likely needed in order to form memories useful for navigation, and demonstrate it using a computational model of the Mushroom Bodies embedded in an agent moving in through a virtual 3D world.

## Introduction

In the insect realm, hymenopteran central-place foragers such as ants, wasps or bees, spend most of their life in the nest, but there comes a time where they are faced with a great challenge for their survival and the survival of the colony: they must wander outside in search for food, and be able to bring it back home safely. Navigating alone through the complex world that lays beyond the bounds of their nest is no easy task, but despite a very small brain, these animals display remarkable abilities to do so using vision.

Naive foragers venturing outside for the first time typically spend 2 to 3 days [24, 45] performing very stereotyped sequences of movements around the nest entrance called learning walks in ants (or learning flights in bees and wasps). This serves to gather the necessary visual memories that will later be used for visual homing [71, 99, 70, 89, 24, 25, 26, 28, 12, 45, 92, 68]. In addition to the learning of these nest-centred views, ants and bees also visually learn and follow idiosyncratic routes within their foraging territory [49, 56, 44, 59].

Even if it is still not completely clear how these views are learnt, stored, or used for an actual navigation task, the advancing progress in the description of the insect neural circuits start to shine light on the matter. Indeed, there is now direct evidence that the Mushroom Bodies (MB), a pair of prominent and evolutionary conserved neuropils, are required for view-based navigation in ants [47]. More specifically, impaired MBs cause defects in the use of learnt views but not in the innate response to visual cues [8], suggesting an important role of the MBs specifically in learning and memory for navigation.

How the MBs’ neural circuits can encode memory has been mainly studied in the context of olfactory learning, in flies and bees [75, 11, 94, 34], but way less in the context of visual learning [50, 90, 20]. The main characteristic of the MBs is the sparse encoding happening there [75], across the thousands of intrinsic Kenyon Cells neurons (named after Kenyon [48]). The axons of these Kenyon Cells represent the bulk of the MB structure, while their dendrites compose the MBs’ Calyces – the main input region [35, 34]; their terminal axons compose the MB’s lobes – the output regions. The Calyces of the MBs can be segregated in three subregions receiving respectively olfactory input, visual input, or both [64, 30, 23].

Interestingly, the volume of the MBs and of the visual input areas in the Calyces has drastically increased in Hymenoptera that have a nest where they need to come back to, suggesting that the ability to store complex visual memories may have allowed the evolution of central-place foraging in these species [81, 22]. Multiple lines of evidence, such as in *Cataglyphis* ants [32], show the existence of optical tracts (optical calycal tract, OCT; anterior superior optic tract, ASOT) projecting from the Medulla and Lamina of the Optic Lobes to the Calyces of the MBs. Such massive visual projections to the MB appear specific to hymenopterans [18, 63, 32].

Remarkably, the MB circuitry modelled at first [34] for olfactory learning turns out to be very suited to also encode visual memories for navigation [3, 97, 27, 102]. Notably, Ardin et al. [3] showed that a bio-plausible MB model of olfactory learning [100] could be directly used, by replacing the olfactory projections by visual ones, to learn multiple images along routes in a virtual environment and to distinguish previously seen images from novel ones. In this model, visual information is sent directly to the MBs – this contrasts with reality. Indeed, contrary to the single relay observed in olfactory projections, visual information in insects transits through several neural relays before reaching the MBs, notably in the Optic Lobes (OL), which are themselves composed of relays in the Lamina, the Medulla [18, 32] and optionally the Lobula and Lobula plate [55]. Olfactory worlds and visual worlds are inherently different in their spatial properties and their task relevance, so this difference is perhaps not too surprising.

The exact nature of the different types of processing happening along the visual relays is still unclear, and how these could actually serve (or harm) navigational performance is entirely unknown.

Our aim is here to use a computational approach to investigate how biologically relevant visual processing can impact the encoding and retrieval of visual memories in the MB. We present a MB-inspired bio-plausible model, but with added biological constraints upstream of the MBs. Specifically, we explore the effect of known mechanisms such as colour opponency [65]; lateral inhibition between neighbouring ommatidia in the lamina and medulla [88, 43]; the pruning of Projection Neurons to KC connections known to happen upon the beginning of foraging life [82, 9, 29]; and the input connectivity and sparsening activity of the KCs [58].

Overall, this works show how these added processing steps are key for the learning of long routes in a complex 3D environment and allow efficient visual navigation at very little computational cost.

## 1 Methods

### 1.1 3D Environment

To investigate how the MB circuitry can encode visual memory for navigation, a virtual environment accurately reproducing visual characteristics of real natural scenes was needed. To this end, we used a dataset generated by the software Habitat3D [80]. This tool provides near-photorealistic meshes of natural scenes, from point clouds acquired with a LiDAR scanner (IMAGER 5006i, Zoller+Fröhlich GmbH, in the case of the dataset used here) [93]. The point cloud used here is mapped on a typical natural habitat of ants, represents an area of about 30 meters radius, and features large eucalyptus trees and several smaller vegetation items. This dataset can be found on the Insect Vision webpage (https://insectvision.dlr.de/3d-reconstruction-tools/habitat3d). The original high-resolution mesh was downsampled in Blender [13] and exported in Polygon File Format where each vertex is defined by 6 values(*x, y, z*, Red, Green, Blue). We ran the simulations with a custom OpenGL-based 3D renderer.

For the agent’s displacements, we only consider the horizontal (*x* and *y*) axes, with the third (*z*) axis being calculated at all times by vertical projection onto the mesh under the agent. In other words, the viewpoint follows the ground topography as if it hovers it at a constant height. We only consider here changes in yaw, not pitch or roll. The impact of latter two on the use of views could be considered in a future study. Non-ground elements (i.e., vegetation) are walk-through. The sky has no texture in order to facilitate the distinction between sky and ground (as can be achieved by navigating insect using the UV-green channels opponency [38] and has been replaced by a uniform blue colour (see section *Colour processing*).

### Insect eye model

Visual input from the virtual world is acquired through a 3D model of the insect faceted eye. This faceted eye model is generated via subdivision of an icosahedron, where the latitude and longitude of each vertex correspond to the viewing direction of an ommatidium – or one Lamina cartridge (in other words, each ommatidium can be considered as one face of the associated dual polyhedron).

In this work, just like the crystalline cones of the ommatidia focus photons from a particular viewing angle (“*acceptance angle*”) onto the rhabdoms [41, 96], each modelled ommatidium covers an acceptance angle and pools all pixels from the given region. We set here the acceptance angle of all ommatidia to approximate the interommatidial angle, meaning that we sample the full visual space. In other words, the acceptance angle is inversely proportional to the number of ommatidia (i.e., to the number of icosahedron subdivisions). We fixed the number of ommatidia for all experiments to 162 (this corresponds to 5 subdivisions of an icosahedron). Consequently, each ommatidia covers a viewing angle of 18.0^*°*^ and takes as input the three colour channels of the OpenGL-rendered 3D environment.

### Insect brain model

Similarly to the other typical MB models, the ommaditia/cartridges (see *Colour processing* section) act as the input space, the Mushroom Body intrinsic neurons (KCs) as the high-dimensional space, and the Mushroom Body Output Neuron (MBON) as a linear classifier (fig. 1).

**Figure 1.**
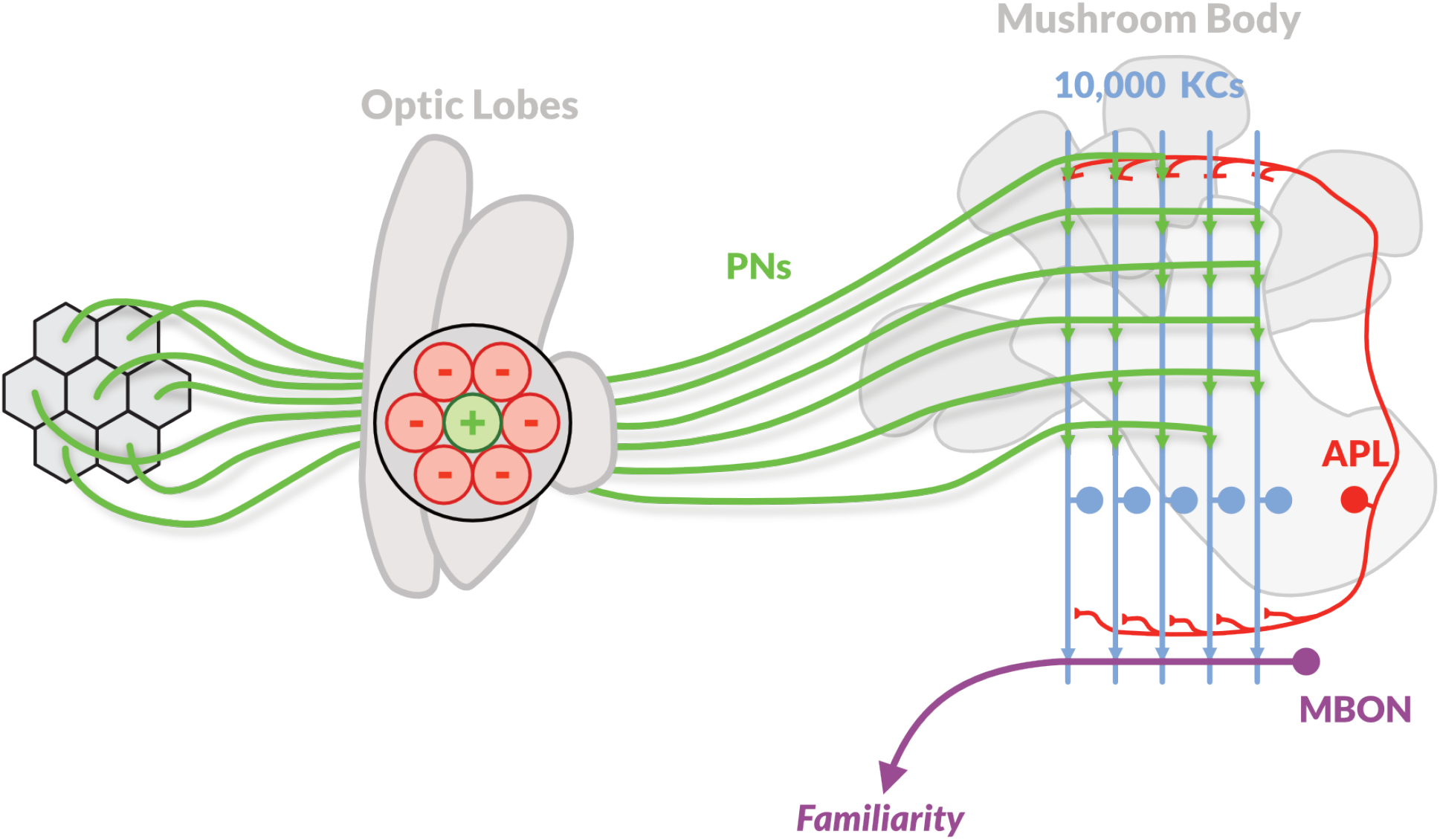
Model overview. Left, seven of the 162 ommatidia; Middle, an example of the centre-surround inhibition across each group of seven ommatidia; Right, An example of how the fan-out connectivity happens in the Mushroom Bodies: each Projection Neuron (PN) connects to a small number of random Kenyon Cells (KCs) in a large pool of 10,000. On average, each PN connects to roughly the same number of KCs, and each KC receives input from the same number of PNs. The Anterior-Paired Lateral (APL) neuron receives excitatory input from all the KCs and in turn inhibits them all, acting as a negative feedback loop which selects only the highest firing KCs at any time. The KCs synapse on Mushroom Body Output Neurons (MBON), after learning (i.e., modulation of the KC-to-MBON synapses), the activity of an MBON encodes the familiarity of a given PN input pattern.

#### Neurons activation functions and Synaptic weights

As our aim in the present work is not to model the neural response dynamics at a cellular level, but rather to investigate the interplay between multiple larger-scale biological processes, we chose not to model the membrane potential nor the sinusoidal response of our neurons. Instead, the PNs are encoded as simple firing rates ([0.0, 1.0] interval), with additive integration for the excitatory postsynaptic potentials (ePSP) in the KCs dendrites. The KCs’ firing activity is encoded as binary values (logical 0 or 1), rather than continuous. Indeed, KCs are known to fire only rarely and timely: usually one action potential at a key timing during the Local Field Potential (LFP) oscillatory rhythm [51], in essence acting akin to a binary population code.

Synaptic weights in our model are fixed, with the exception of the two layers where some form of learning may happen: the PN-to-KC synapses, and the KC-to-MBON synapses. In both cases, these synapses follow an instantaneous weight change (initialised at +1.0, fully excitatory; set by learning in one shot to 0.0). This simplification enables us to ensure the effects observed are not dependant on a gradation of synaptic weights, where models can be infinitely accurate (beyond biological reason).

#### Colour processing

Both in the retina and the optic lobe, a columnar organisation is preserved (retinotopy). In the Lamina, these columnar units are often called “cartridges”. Each cartridge receives input from the central receptors of the overlying ommatidium and from the surrounding receptors of neighbouring ommatidia. Thus, a given cartridge receives input from a small group of ommatidia that are all pointing towards the same direction in the visual field [61, 40, 78, 77, 79].

As in seemingly most ant species, we assume that our agent has two receptor cell types which respectively detect UV and Green light [67, 1]. Assuming that the sky’s global irradiance represents naturally the major source of UV light [10], we render in our simulation the sky as uniformly blue and use the blue and green channels from the render to activate the agent’s UV and green receptors respectively. Thus, the first component of our model converts the RGB values from the 3D rendering that each ommatidia receives, into a single output firing-rate, therefore approximating both the non-opponent achromatic luminance detection in the Lamina and the colour-opponent (subtractive) detection in the Medulla [66, 85, 36]. We look at two different UV-Green colour opponency, for Green dominency (eq. (1), left), or the UV dominency (eq. (1), right):

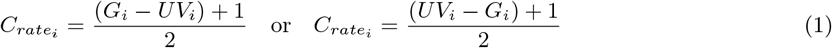

where *G*_*i*_ is the green receptor value of ommatidia *i* (and *UV*_*i*_ for UV receptor *i*), *C*_*rate*_ is the single firing rate value of cartridge *i*. Note that we refer to this input space using *ommatidia* or *cartridges* terms indifferently in this work.

#### Lateral Inhibition

Lateral inhibition is very common in the sensory systems [88], and is usually assumed to be implemented in a temporal fashion in the Lamina, using the delay in the co-incidence of a stimulus between two adjacent receptors [43]. Examples of biological applications of it include orientation selectivity, and feature detection [7]. Here, we circumvent the non-linearity that is typically associated with such visual processing [98] and directly implement our lateral inhibition like so:

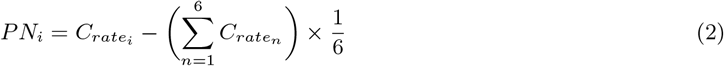

where *PN*_*i*_ is the total input to the PN *i* and 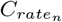 are the firing rates of the *n* = 6 neighbouring cartridges for cartridge *i*.

#### Projection Neurons layer

The output of each cartridge is then conveyed by a layer of visual Projection Neurons (PN) to the MB. This layer is composed of as many neurons as there are of cartridges. Before learning, the output synaptic weights (onto KCs) are all set to +1.0, so the output of this layer onto the KC dendrites is directly proportional to the excitation of the PNs for any given view.

#### Kenyon Cells layer

The Mushroom Body model is similar in its overall structure as the one in Ardin et al. [3]: this layer is composed of a rather large number of Kenyon Cells, set for most of the experiments (unless otherwise indicated) to 10,000 KCs. This represents an underestimation compared to real ants [32], but enables fast computation times, while remaining conservative about memory space. The PN-to-KC connectivity pattern follows a pseudo-random rule where each KC receives a fixed number of inputs from randomly chosen PNs if and only if their output synaptic weights are not 0.0. Conversely, all PNs will connect in average to the same number of KCs. This connectivity scheme provides the fan-out pattern that is essential to the sparse encoding [51, 5]. Following the additive integration approach, each KC’s dentritic ePSP will correspond to the sum of the inputs it receives: all the PNs activity that synapse onto the KC. This would normally require re-normalisation of the values to fall into the [0.0, 1.0] range with help of an activation function, however this is not needed here: whether a KC will fire an action potential or not (0.0 or 1.0) depends on the population activity of KCs and on the sparsening activity of the APL neuron (see **APL-like normalisation** paragraph).

#### Projection Neurons pruning

We add an approximated learning walk in the virtual environment, during which views are acquired from random positions contained in a circular area around a fictive nest. Then, we take the standard deviation of the activity for each PN across these learning walk views and compare it to a threshold: this threshold is the mean of the standard deviation values for all PNs. Every PN which falls under threshold is tagged for pruning. Then these PNs are eliminated, and the pseudo-random connections are regenerated, approximating what happens biologically (see **Discussion**).

#### APL-like normalisation and sparse encoding layer

We model the APL-neuron equivalent (called APL in drosophila [58], A3-v in honeybees [14, 83]), which acts as a filter: this neuron applies an inhibitory feedback to the whole KC population, of a strength that is proportional to the sum of the inputs coming from that same KC population [58]. This results in a *winner-takes-all* selection of the most active KCs for production of a spike. We implemented this normalisation in a relative fashion, where a defined portion of the KCs with the highest ePSPs get selected. The selected firing KCs can thus be coded as a binary matrix (1 = spike, 0 = no spike).

#### KC-to-MBON plasticity

Prior to any learning, all the KC-to-MBON synapses are set to their maximum value (+1.0), so that given additive integration, the firing rate of the MBON is equal to the proportion of selected KCs. The output value of the MBON therefore can be interpreted as representing an *unfamiliarity value* for the current view, which is at first equal to 1.0. In other words, prior to learning (and as expected), any scene appears totally unfamiliar to the model.

During learning, the output synapses of the active KCs are depressed (their weight goes from +1.0 to 0.0). Thus, after learning, if a familiar scene is presented again, the same KCs will fire again, but their synapses to the MBON will have been set to 0.0, meaning that the output of the MBON will be 0.0, that is, the view will appear familiar. Any view that will be somewhat similar to a learnt one, even not identical, may thus activate a similar pattern of KCs and thus produce a familiarity value that will vary between 0.0 and 1.0.

## Results

In order to characterise the effect of known visual pre-processing (lateral inhibition between signals of neighbouring ommatidia, pruning of the projection neurons) on the encoding and use of view for navigation, we look at the change in activity of the MB input layer (PNs activity) (fig. 2) and the model’s familiarity output (MBON activity) along elementary displacements of the agent in the virtual world (fig. 3). These elementary displacements are of two types: either rotational, where we look at the change in neuronal activity between a reference position and a full 360^*°*^ rotation on the spot, taking a view every 5^*°*^(fig. 2a, fig. 2c, fig. 3); or translational, where we look at the change in neuronal activity between a reference position and a set of 40 increasingly distant positions (on a log space) in a random direction (fig. 2b, fig. 2d, fig. 3). For this experiment, the MB model is set with 10,000 KCs; each KC receives input from 25 random PNs; and the APL normalisation enables 1% of the most excited KCs to fire an action potential. But as we will see, the insight resulting from this investigation is independent of these parameters values.

**Figure 2.**
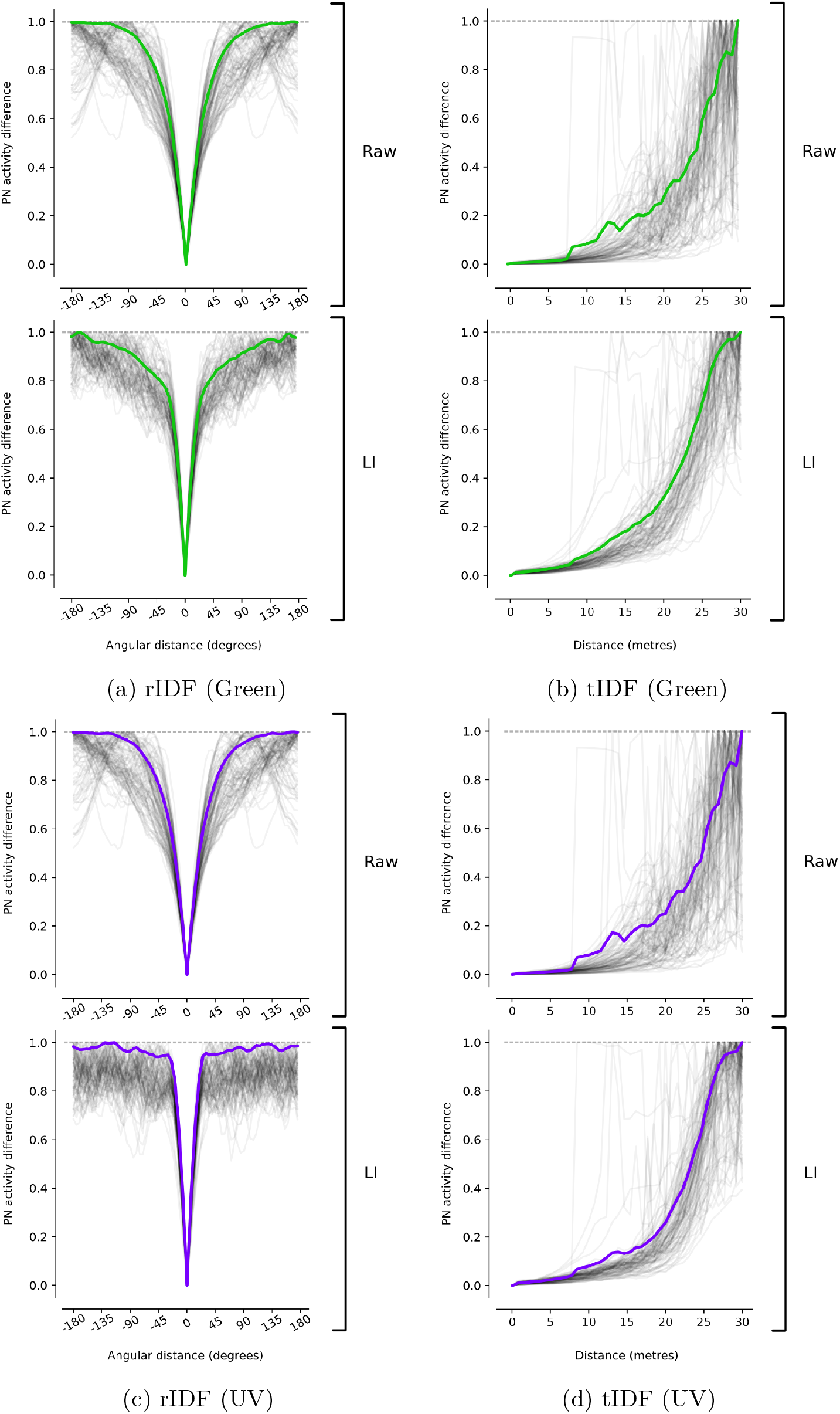
PNs Image Difference Functions. Difference between a reference view and the other views along either a rotation in place (*a, c*) or a linear displacement (*b, d*). *Raw*, raw image; *LI*, With lateral inhibition. *a, b*, Green dominancy (*Green*-*UV*); *c, d*, UV dominancy (*UV*-*Green*).

**Figure 3.**
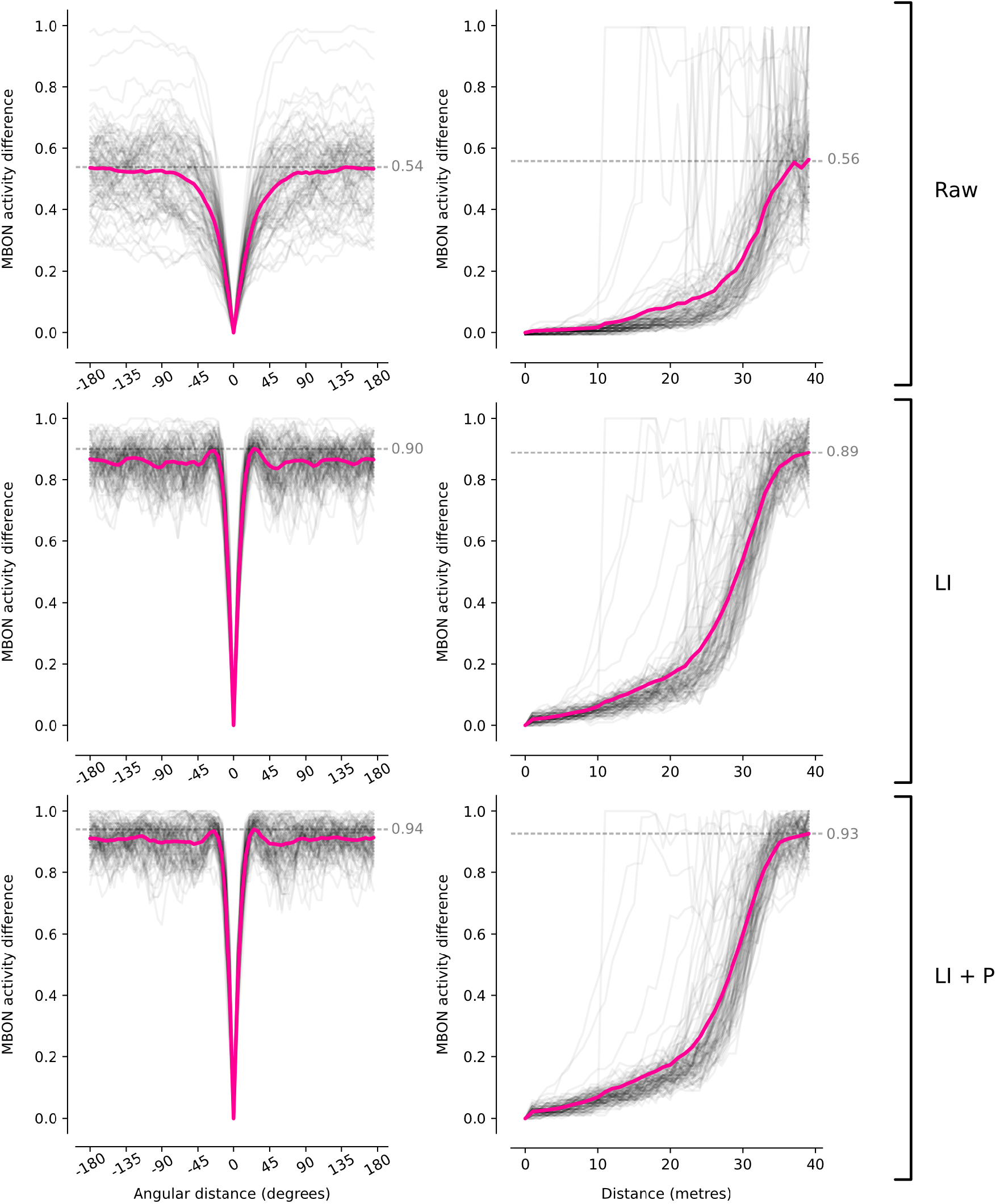
MBON Difference Functions. Difference between a reference view and the other views along either a rotation in place (*left*) or a linear displacement (*right*). *Raw*, raw image; *LI*, With lateral inhibition; *LI + P*, With lateral inhibition and after pruning.

First, in the input layer (PNs) (fig. 2), we look at the difference in the response profiles in absence and in presence of lateral inhibition between the signals of neighbouring cartridges: without lateral inhibition, the PN layer response follows a rather classical rIDF profile [91, 106], with some amount of similarity extending up to 160^*°*^ around the reference position (fig. 2a, fig. 2c). With lateral inhibition, we observe a sharpening of the rotational selectivity of the response: the PN layer’s response shows similarity for only 60^*°*^ around the reference position (30^*°*^ on both sides). This effect is not surprising, as the centre-surround antagonism provided by lateral inhibition is known to increase the ability to discriminate [60, 101]. Results are similar for the translational movement (fig. 2b, fig. 2d): lateral inhibition sharpens discrimination in the sense that the similarity of the PN layer’s response changes more quickly across displacement and reaches the maximum discrimination quicker than without lateral inhibition.

But it is by looking at the MBON familiarity output (fig. 3) that we can observe the key importance of early lateral inhibition. Without lateral inhibition, the maximum MBON activity stays around 0.5, meaning that, at the MBON level, completely different views (such as the ones 180^*°*^ opposite of the reference one) still appear 50% similar to the reference one. This value drastically improves with the presence of lateral inhibition: the model can fully differentiate between the reference view and other views. This observation is also true for the translational experiment, where the presence of lateral inhibition bumps the discrimination ability from 56% to 89% after 35 m. In a sense, lateral inhibition as observed in the Lamina of insects [62, 43, 39] is strongly improving discriminability between familiar and unfamiliar views in the MB. The mechanistic reason will be investigated subsequently.

The addition of the pruning of projection neurons further improves the discrimination range in the MB, but its importance will appear more clearly in the following experiments.

We know that in the MB, only the KCs receiving the strongest inputs (highest ePSP) will fire a spike and transmit information about the current scene further. Therefore, it is important that in the PN layer, the PNs that fire strongly actually carry useful information for scene discrimination. Whether a PN is discriminative or not (that is, whether it carries useful information about the scene’s identity) can be simply quantified by looking at the standard deviation of the PN’s response across displacement along a route: the higher the variation in time, the more information it carries about the identity of the view experienced. Indeed, if a given PN’s activity remains constant across displacements, it carries by essence no useful information for recognising locations. We therefore observed the variation of activity of the PNs along a random walk in the virtual world, in relation to their average activity (fig. 4).

**Figure 4.**
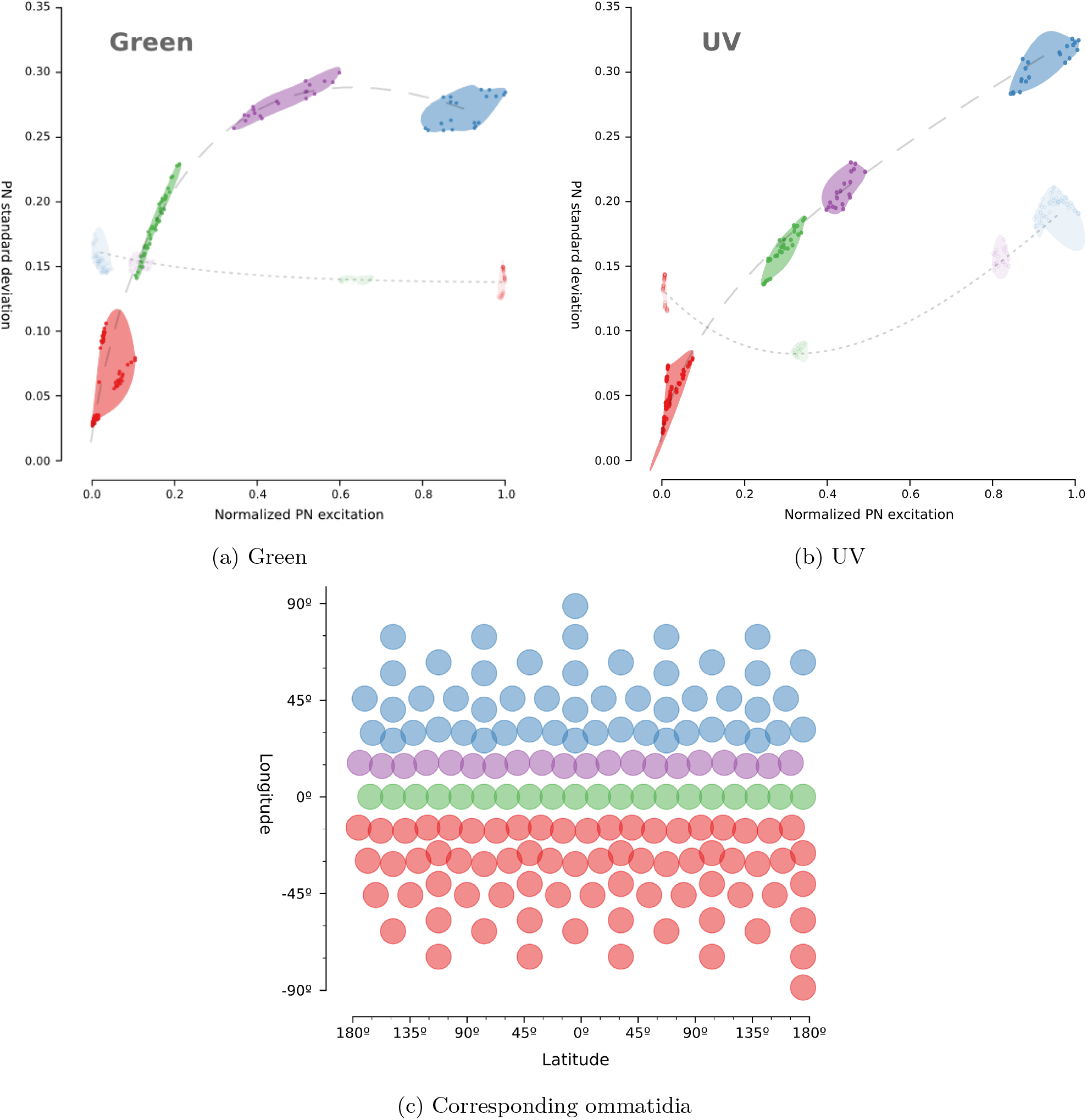
PNs informativity. PNs activity along random routes, in relation with the standard deviation of that activity. A high *sd* means the PN firing profile varies a lot; a low *sd* means the PN rarely fires or always fire. *a*, Green dominancy (*Green*-*UV* opponency); *b*, UV dominancy (*UV*-*Green* opponency); *c*, Corresponding ommatidia directions for each cluster in *a* and *b*. In *a* and *b*, Transparent points and clusters (and dotted line): Raw image; Opaque points and clusters (and dashed line): after Lateral Inhibition. (c) Lateral Inhibition + Pruning

This repartition of the PNs activity shows important clustering of the response profiles. Like in the Medulla, each PN we modelled responds to a specific zone of the visual field, and thus can be associated to a corresponding latitude and longitude in the panoramic scene where the ommatidia points. We can thus identify clusters of PNs responding to the region of the sky; the ground; the upper horizon; the lower horizon (fig. 4c). As expected, the PNs responding to the region above the horizon carry the most information about scene identity, because this is where most of the change in panorama happens due to the relative movement of the trees and bushes. However, we observe that without lateral inhibition, there is no clear relationship between the average and variation in activity of PNs. The most active ones do not necessarily carry information allowing scene recognition (see fig. 4a, fig. 4b). For instance, PNs responding to UV receptors (*UV − Green* opponency) (fig. 4b) are most strongly activated in the region that points to the uniform sky area, meaning they’re uninformative for scene recognition. The problem is even starker when considering the green receptors (*Green − UV* opponency), as seen with the poor vertical separation of the clusters in fig. 4a. Without lateral inhibition, the most active PNs – the ones that will contribute most to the KCs’ activity – are pointing below horizon, that is, a region that carries little information to discriminate scenes reliably. In this virtual world (like in reality) the most informative region corresponds to the above horizon (fig. 4c). The addition of lateral inhibition enhances the vertical separation of the clusters, silencing the least informative PNs that point in visually unchanging regions, and enhancing the response of the informative PNs that detect contrasted edges, which will vary across displacement and thus carry information for discriminating scene.

In our simulations, the opponency between *UV − Green* or *Green UV −* result in very similar information to that of *UV* and *Green* taken alone. This opponency might however be useful to cancel illumination intensity variations across the eye [15], but this was not required here given the constant light intensity in our virtual environment.

We then investigated the effect of the PNs-to-KC connectivity on the efficiency of the KC layer to discriminate across different locations along a long route. We look at the variation in the input profile of the whole KC population across multiple random routes, for all possible number of PNs per KC (i.e., from 1-PN-per-KC, to all-PNs-per-KC) (fig. 5). For each PN-per-KC condition, we run 30 individual random routes of 500 steps, 5 mm per step. The MB model used in this experiment was limited to a number of 1,000 KCs for speed purposes, but this bears no qualitative effect on the results, as the connectivity effect is function of the ratio of PN to PN-to-KCs connections, and 1,000 KCs provide a good enough resolution for this endeavour. For each possible connectivity, we quantify whether the KCs’ inputs profiles along the route tend to correlate, using a Pearson correlation coefficient across all possible pairs of KCs. This enables us to quantify the proportion of these cells that are redundant. Indeed, one expects an optimal system to tend to the lowest number of correlated KCs. Figure 5 shows the mean correlation coefficient across the 1,000 KCs, as well as the total number of KCs that are correlated more than 95% of the time. According to combinatorics and given by a binomial coefficient, the maximum number of possible PN combinations that KCs could randomly sample is reached when each KC samples half of the existing PNs [46]:

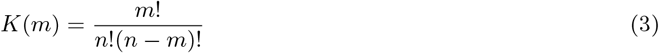

where *K*, the number of possible different PN combinations that KCs receive is maximum for *m* = *n/*2 if each KC samples *m* PNs out of *n*.

**Figure 5.**
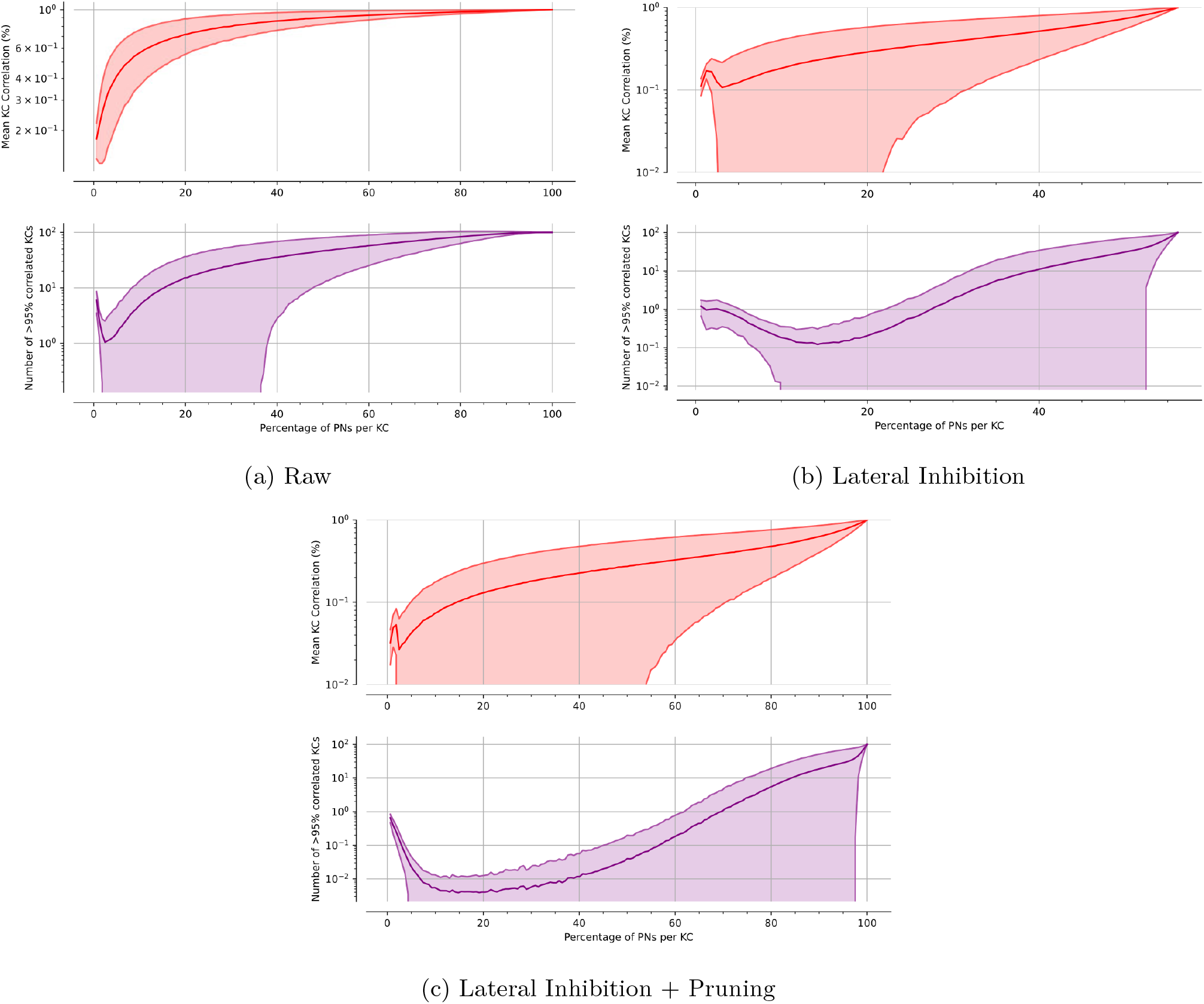
KC layer correlation. Correlation of the KCs firing profiles (10,000 KCs) along random routes (3 m), in relation with the number of PNs-per-KC synapses. *a*, Raw image; *b*, With lateral inhibition; *c*, With lateral inhibition and after pruning. *Red*, mean correlation; *Purple*, Number of KCs correlated more than 95% of the time.

Contrastingly, in our simulation we observe that the optimal ratio of PNs per KCs connections falls to much lower values: between 2% and 5% of the PNs without lateral inhibition (fig. 5a), and between 10% and 20% of the PNs with lateral inhibition (fig. 5b). With the pruning of PNs, the number of KCs firing in correlation rises slightly (particularly visible on the mean correlation curve in fig. 5c, top): this is due to the fact that less PNs overall synapse to the KCs population, diminishing the absolute discrimination power of the system. However, the trade-off in terms of PN per KC connectivity remains the same. The reasons and consequences of these results will be discussed below.

In order to observe how much memory space our MB model could provide, we observed the risk of *aliasing* in the familiarity output as a function of the length of a route learnt (fig. 6). To that end, we made the agent learn progressively longer routes in the environment (from 0 up to 60 metres), while continuously testing its ability to discriminate between familiar and unfamiliar views sampled either at very distant position in the world (‘remote views’) or in a completely differently structured visual world (‘alien world’) (fig. 6). This allows to assess how *mistakenly familiar* novel views appear as the model learns an increasingly longer route. If the memory space were to be completely exhausted by learning (e.g., all KC-to-MBON synapses set to 0.0), any new view, no matter how novel, would appear familiar. Figure 6 shows that with direct ommatidial activity input to the MBs, the unfamiliarity of novel views decreases sharply with the length of the learnt route (irrespective of whether these novel views were taken from the same world or the alien world). For instance, given a memory capacity of 10,000 KCs, novel views appear more than 85% familiar after the model has learnt a 10-metre long route only (fig. 6, grey and red curves). In contrast, with the lateral inhibition added, and after the pruning of PNs-to-KCs connectivity, the novel views can still be identified as unfamiliar (>50% unfamiliar) even after learning a 60 metres route (fig. 6, green and yellow curves).

**Figure 6.**
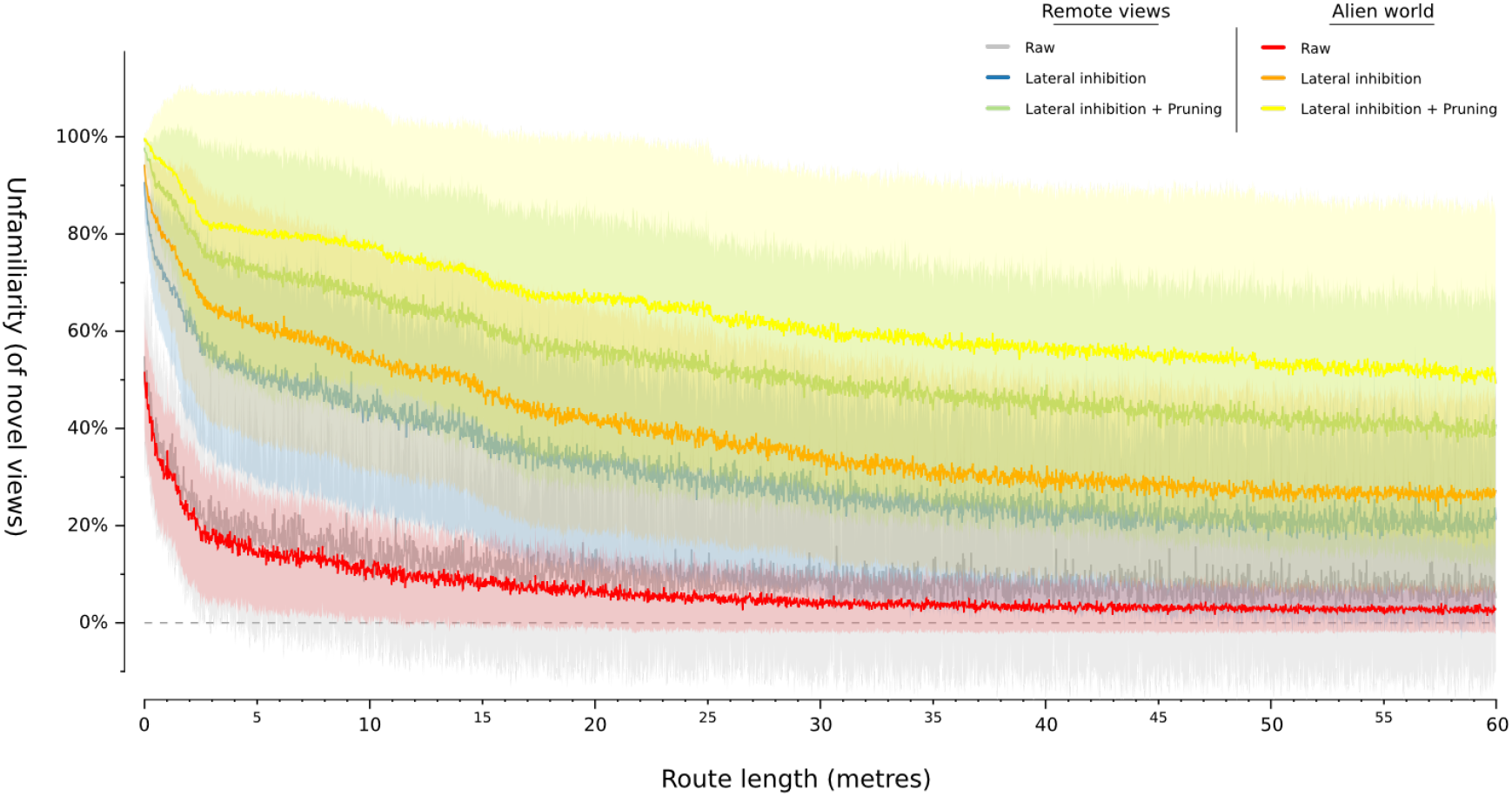
Memory saturation. Apparent unfamiliarity of novel views as the model learns long random routes (60 m). *Remote views*: novel views in the same world; *Alien world*: novel views from a differently structured world.

## Discussion

The visual world is not random: it is highly structured and spatially correlated (even more so along a particular ant’s route). This is contrasting with the olfactory world, and it is thus sensible to assume that the MB would account for this structured nature of the inputs to increase learning efficiency. Our modelling approach reveals here how visual pre-processing, appropriate inputs sampling statistics and simple pruning rules can enable the MB to do just so.

### The MB is not made to accommodate raw ommatidial input

We show that the sparse coding present in the MB circuitry is well suited to encode, store and compare views, and that it allows for navigation in complex environments, similarly to Ardin et al. [3]. However, we show that sending visual information directly from the ommatidia – the ‘raw view’ – to the KC population suffers from severe aliasing, quickly exhausting memory space. This problem does not result from an absence of information in the ‘raw view’, but instead arises in the transfer from the ommatidial input layer to the high dimensionality code in the KCs layer.

### Lateral inhibition ensures PN activity carries information

As only the KCs that receive strong PN inputs are likely to fire, a desired outcome would be that the PNs that fire strongly (and thus excite the KCs the most) should be the ones to carry navigation-relevant information. This is not the case in the raw view, where the PNs corresponding to the UV-intensive sky (high up above horizon) and green-intensive ground (below horizon) are the ones that fire the most. These two regions present few variations across different locations, and thus many KCs fire similarly across most locations – they are therefore useless for navigation.

Remarkably, this problem is simply solved with the addition of lateral inhibition in the input of the PN layer. This process is well-known in early visual processing in the Lamina [60, 43, 39]. The usual observation of the role of lateral inhibition in compound eyes is multiple. For example, it can improve the angular sensitivity of the eye (i.e., it sharpens the image) [60], allowing better discrimination of the views, as seen in the rIDF profiles. Or it can act as an elementary filter for edge extraction [39] (akin to a Haar-like feature detector [31], which are classically used in machine vision applications [95]). It can also serve for feature (target) extraction or movement perception [7].

These roles of the lateral inhibition may indeed be advantageous for image processing in general, but the benefits we aim to highlight here are of a different nature and directly depend on constraints in the ways MB function. Early lateral inhibition does not so much improve the amount of information available in the PN layer (fig. 4) but enables to adapt the code mediated by the PN layer to ensure that this information will be carried out in the downstream stage of the KCs. Only the most stimulated KCs will fire an action potential, therefore, it is of great importance that the PNs that carries information for discriminating a scene at a given time are the ones that fire the most. By ensuring that PNs respond to the presence of contrasted edges in the receptive field, lateral inhibition ensures that the strongly firing PNs carry information that necessarily changes across locations – information that is therefore relevant for navigation. This in turns enables the KC layer to convey relevant information as well, making the MBON classifier to be fully exploitable.

In the absence of lateral inhibition, many KCs fire in response to inputs from PNs stimulated by non-informative regions of the eyes. Thus, the effective pool of useful KCs is strongly reduced, and memory load is lowered. Impressively, the simple addition of lateral inhibition enables a huge improvement (fig. 6) of the length of the route that the agent can learn before saturating its own memory.

### Pruning of PN-KC connections during early learning walks improves the system

However, with lateral inhibition only, some KCs still appear to fire in an uninformative fashion: either they fire most of the time, or never (fig. 4). Even if PNs respond to the presence of contrasted edges, due to the random connectivity, some KCs may end up receiving exclusively high-firing PNs, or rarely firing PNs, making them all that much uninformative as well. Reorganising this random connectivity can solve this problem, and that is exactly what can be seen in these insects.

It has been shown that the transition associated with the polyethism from nest workers to foragers in *Cataglyphis* ants resulted in important neuronal plasticity in the visual input synaptic complexes (micro-glomeruli) of the MB Calyx [87, 90]. This neuronal plasticity corresponds to a reduction of the micro-glomeruli density: an increase of the overall Calyx volume associated with a decrease in the number of micro-glomeruli. This decrease in density of (presynaptic) visual projection neurons is associated with a dendritic outgrowth of the (postsynaptic) Kenyon Cells in the MB Calyx [87, 90]. In other words, the onset of foraging triggers the axonal pruning of the visual projection neurons. Importantly, this is independent of the ant’s age and requires several days of light exposure, suggesting that these changes happen during the early Learning Walks period displayed by these ants, and are helpful for preparing the visual pathways to the impending navigational tasks [89].

Remarkably, we show that adding a simple pruning rule (see **Projection Neurons pruning** paragraph) after a learning walk does effectively further improve the effective memory capacity. Literally, this pruning can be viewed as an adaptation of the eyes to the type of visual information that is informative in the specific environment where the insect has to navigate in. The learning walks (that is, the movement) seem essential to assess *what* is informative. This is exploited by our simple pruning rule by looking at which PNs vary and which do not. How this rule can be implemented biologically remains to be seen, but we can easily assume that KCs lose their connectivity to PNs that have no change in their firing activity [82, 9].

### Visual worlds are different from olfactory worlds: impact on the MB connectivity

For maximising the separation of inputs in the high dimensionality KC layer, combinatorics indicate that each KC should connect to half the PN population [46]. For our 162 PNs, this means there are 3.66×10^47^ connectivity patterns (see eq. (3)). This theoretical prediction may very well be useful in the case of olfaction, and actually was verified in the Locust olfactory system. However, we see here that this does not equally apply to visual information, as many KCs are similar to other KCs in their firing profiles along routes. This is because the visual world is strongly spatially structured and visual information varies drastically across the vertical axis of the eye regions in a way that is quite constant over multiple locations. Thus, if all olfactory receptors may have an equal chance of being useful, this is not the case with visual receptors given a navigational task. Our analysis shows that to avoid redundancy, KCs should connect in average to only 5-10% of PNs. Even if the theory tell us that connecting to more PNs (around 50% of PNs) would increase the number of permutations and thus the likeliness that the connectivity pattern of each KCs is unique, this does not necessarily apply to their input across a route. Note also that connecting to 5% of 162 PNs is already sufficient to produce 9.87×10^12^ different possible connectivity patterns (see eq. (3)), which given the tens (or hundreds) of thousands of KCs present in ants, more than enough to ensure that the probability of KCs having the same connectivity pattern would be extremely low.

We could thus draw as prediction that the Calyces of the Mushroom Bodies of visually navigating insects such as bees and ants should show less diverse connection profiles in the collar (visual) than in the lip (olfactory) areas of the Calyces.

### Continuous learning during navigation is not a problem, perhaps a solution

Early models of visual navigation often encountered the problem of the number of views stored, and thus needed a mechanism to decide *when* to learn a view. This was a difficult task given that in natural environments, the view changes smoothly with displacement [106]. We show here that, by learning the view at every step of the agent (every 5 mm) and given the size of the virtual world (route across 60 metres), our model essentially achieves continuous learning. This was also the case in the previous (albeit non based on the MB circuits) model of Baddeley et al. [6]. Over-sampling is not a problem here, given the fact that if the view is already familiar (e.g., it has been learnt 5 mm before) likely the same KCs will fire, and no change in connectivity in the KC-to-MBON synapses will happen. In other words, learning is only triggered by novelty. Despite such continuous learning, we can see that 10,000 KCs only are sufficient to memorise a 60-metre long route without memory saturation (fig. 6).

### Potential improvements to the MB model

Two main classes of KCs have been described. Class I KCs receive input from numerous single spine PNs, and consequently require coincident inputs in order to be depolarised. Class II KCs however receive most of their inputs from a single PN, and thus convey more specific information [21, 29, 19]. Our model only implements the former, which is also by far the most common type of KCs in hymenopterans [21]. Class II pathways enable the detection of innately relevant specific stimuli in the context of olfaction (such as food or pheromonal odours [107]). Their role in the context of vision could be to signal the location of innately significant visual cues, such as trees for arboreal ants, or flowers for bees, but this remains to be investigated.

Also, we implemented here only one MBON for linear classification of the view familiarity, while insects Mushroom Bodies possess many such output neurons. In the context of olfaction, the different MBONs can enable the association of odours with different valences such as aversive/appetitive, or different needs such as water, sugar or amino-acids [4, 37, 73, 84, 76]. The same is true in the context of vision, where different MBONs can convey signals for attraction/repulsion or left/right opponent pathways, which prove to be key for visual navigation ([104, 52, 69, 103]). In any case, our current conclusions about the impact of visual pre-processing on memory efficiency should benefit all these parallel output pathways equally.

Furthermore, there is evidence in bees of learning happening in the Calyces of the MBs, where a large octopaminergic neuron (*V UM*^*mx*1^) encodes the innate response to sucrose and modulates the KC sensitivity to the PN inputs [33]. We can see how the potentiation of the PN-to-KC connection strength for important locations in the world (i.e., specific places previously associated with food or negative events) would increase the chance to trigger these KCs, and thus lower the threshold for the recognition of these particular locations.

Regarding learning dynamics, it is worth noting that we modelled here learning as a simple, immediate and full depression of the KC-to-MBON synaptic weight (from 1.0 to 0.0 given a single KC spike), but the implementation of a smoother synaptic weight change would mean that only the KCs that fire in a stable way across several steps would contribute to learning. Thus, this should allow the model to learn distal features of the panorama (which are more useful) quicker, and filter out the proximal noise (e.g., small blades of grass that pass by the retina quickly). However, the detection of the different motion patterns between proximal and distal features could also be done on the earlier stages of the visual pathways, as was for instance shown to be the case in lobula plate tangential cells in flies [53].

Another element that would be worth exploring in future models is the role of KC-KC excitatory connections or gap junctions [54, 57]. These could act as a temporal priming of the KC layer response between successive views [72], and thus potentially account for the ability of navigating ants to learn the sequences of views encountered along their route.[105, 86]. Finally, all projection neurons modelled here sample a small portion of the visual field. It would be interesting to explore the potential role of other PNs types which sample larger areas of the compound eye, such as some neurons taking input in the Lobula [17, 74]. These could potentially convey higher-level image components or frequencies extraction such as rotation-invariant features which would provide robustness to head pitch variation [2]. In the same vein, motion sensitive neurons [42] if projecting to the MB, could enable the learning of dynamic cues such as scene-specific optic flow variations, depth information or occluded edges as evidenced in honeybees [16].

## Conclusion

Overall, our modelling work shows how the storage of visual information in the Mushroom Body can drastically improve given a sensory processing adapted to the visual world proprieties. We show that pre-processing such as lateral inhibition between neighbouring cartridges, appropriate input statistics, and neural pruning at the onset of foraging, can enable a small MB constituted of only 10,000 KCs to ensure the storage of views learnt continuously over more than 60-metres long routes in realistic virtual reconstruction of complex natural environments. This represents a more than 50-fold improvement compared to a MB receiving raw sensory input. Given the hundreds of thousands of KCs observed in the Mushroom Bodies of expert navigators such as ants and bees [64], as well as the myriad of additional visual processing that remain to be investigated, we can start to picture how these insects can navigate so robustly despite their tiny brain.

## Acknowledgements

This research was funded by the European Research Council (grant reference number: EMERG-ANT 759817, author: A.W.)

